# In vivo reprogramming of *Caenorhabditis elegans* leads to heterogeneous effects on lifespan

**DOI:** 10.1101/2024.05.03.592330

**Authors:** Nibrasul Kamaludeen, Yann Mauge, Sonia El Mouridi, Sara Picó, Alba Vílchez-Acosta, João Agostinho de Sousa, Marie Pierron, Viviane Praz, Ferdinand von Meyenn, Christian Frøkjær-Jensen, Alejandro Ocampo

## Abstract

In the last decade, cellular reprogramming of fully differentiated cells to pluripotent stem cells has become of great interest. Importantly, cellular reprogramming by expression of Oct4, Sox2, Klf4, and cMyc (OSKM) can ameliorate age-associated phenotypes in multiple tissues and extend lifespan in progeroid and aged wild-type mice. Surprisingly, the effects of in vivo reprogramming have not been deeply investigated in any other model organisms. Here, for the first time, we induce in vivo reprogramming in *C. elegans* using a heat-inducible system at multiple developmental and adult stages. Similar to mice, expression of the reprogramming factors leads to premature death with different levels of toxicity at distinct developmental stages and aging. In vivo reprogramming in *C. elegans* might represent a valuable tool to improve our understanding of development and in vivo reprogramming.

## INTRODUCTION

Reprogramming somatic cells into pluripotency by forced expression of the Yamanaka factors *Oct4*, *Sox2*, *Klf4*, and *c-Myc* (OSKM) has led to exciting developments of high therapeutic interest^1–3^. Importantly, these reprogramming factors have been shown to act at the epigenetic level by reversing the epigenetic landscape towards embryonic and younger states^4^. Consequently, multiple studies in rodents and human cells have used these or other factors to reprogram differentiated cells towards new cell fates^5^, recover stemness, and slow or reverse age-associated phenotypes both in vitro and in vivo^2,4,6–9^. In mice, continuous induction of in vivo reprogramming leads to organ dysfunction, teratoma formation, and premature death. Importantly, these effects can be avoided by partial reprogramming using short-term cyclic expression of OSKM, resulting in the reversal of age-associated phenotypes in multiple organs and extension of lifespan in progeroid and aged wildtype mice^2,10–12^. However, in vivo reprogramming is still associated with adverse effects such as organ dysfunction, teratoma formation, and premature death^7^.

To date, most reprogramming studies have been performed in human cell culture or in mice^2^. However, induction of in vivo reprogramming has never been tested in *C. elegans*. In this line, in vivo studies in mammals possess multiple disadvantages, such as a longer lifespan, high costs, ethical constraints, and complexity. On the other hand, *C. elegans* has a short mean lifespan of only three weeks when cultured at 20°C. *C. elegans* is a widely used model organism that is particularly suited for epigenetic and cell fate studies due to multiple advantages, including fully identified cell lineages, amenability for genetic screens, small size, and transparency, allowing microscopy acquisitions of whole organisms^13^. In addition to being one of the most commonly used model organism for aging studies, *C. elegans* also provides the benefits of cost efficiency, lower organismal complexity, and conservation of many signaling pathways^14^.

Regarding the feasibility of inducing in vivo reprogramming on *C. elegans*, several genes share a certain degree of homology with the reprogramming factors in mammals^15–22^. Although the induction of in vivo reprogramming has never been described in *C. elegans,* natural trans-differentiation events and ectopic expression of tissue-specific transcription factors in embryos have been reported^17–20^. Moreover, during natural trans-differentiation events, members of the NODE (Nanog and Oct4 associated deacetylase) complex and SOX2 promote the initiation of a natural cellular reprogramming event where a fully differentiated rectal cell transdifferentiates into a neuronal cell^15^. In addition, two more natural trans-differentiation events during development have also been reported in *C. elegans*^20,23^. Regarding the induction of direct reprogramming, ectopic expression of a single *C. elegans* transcription factor has been shown to directly convert mitotic germ cells into specific neuronal types^24^. In addition, another study showed that brief expression of a single transcription factor (ELT-7 GATA) can convert fully differentiated, highly specialized non-endodermal cells of the pharynx into fully differentiated intestinal cells in adult *C. elegans*^25^. All these findings suggest a certain degree of cellular plasticity and collectively propose *C. elegans* as a valuable model to study in vivo reprogramming.

Toward the goal of inducing in vivo reprogramming in *C. elegans*, we first conducted a bioinformatic analysis to identify orthologs of the most commonly used transcription factors to induce cellular reprogramming, including *Oct4*, *Sox2*, *Klf4*, *c-Myc*, *Lin28*, and *Nanog* (OSKMLN)^10^. Subsequently, we selected the *C. elegans* orthologs of OSKL and cloned them into vectors under the control of a heat-inducible promoter to generate transgenic animals with multi-copy arrays to reprogram worms. After optimizing the induction protocol, we characterized the effect of in vivo reprogramming at different stages of *C. elegans* development and aging. Surprisingly, the induction of in vivo reprogramming resulted in aberrant phenotypes such as developmental abnormality, bagging, and premature death. These findings suggest *C. elegans* as a promising model to study in vivo reprogramming and its effects on development, epigenetics, and aging.

## RESULTS

### Generation of heat-inducible reprogrammable worms and optimization of induction protocol

To generate *C. elegans* reprogramming strains, we first identified the orthologs of mouse reprogramming factors Oct4, Sox2, Klf4, and cMyc (OSKM). The four *C. elegans* orthologs, *ceh-6*, *sox-2*, *klf-1*, and *lin-28,* were cloned into individual plasmids and microinjected as a pool of plasmids. After recombination, heritable extrachromosomal arrays containing multiple copies of the reprogramming factors were generated, and reprogrammable worm strains (4F) were selected (Figure 1A and Figure S1A). As the transgenes were under the control of a heat shock promoter (Figure S1A), we first optimized the conditions for the induction and assessed the effects of heat shock on *C. elegans* lifespan^26^. Towards this goal, we tested the effect of different induction temperatures and durations by analyzing *ceh-6* and *sox-2* mRNA expression levels. Induction at 33°C for 3 hours led to higher expression levels of the *ceh-6* and *sox-2* (Figure 1B and Figure 1C). Next, following this protocol, we analyzed the duration of expression over time and observed a peak of expression 4 hours post-induction, which completely subsided within 24 hours (Figure 1D). Based on these observations, we selected this protocol and analyzed the expression levels of all the reprogramming factors 4 hours after heat shock. Importantly, we detected significant levels of expression of all four reprogramming factors in 4F induced worms compared to the control and 4F uninduced worms (Figure 1E). Similar results were obtained by analysis of bulk RNA-seq, where expression levels were higher 4 hours post-induction and decreased after 48- and 72-hours post-induction (Figure S1B). In agreement with these observations, GFP reporter expression was also detected throughout the body upon heat shock (Figure 1F). These observations suggest the successful generation of 4F *C. elegans* and optimization of the induction protocol.

**Figure 1:**
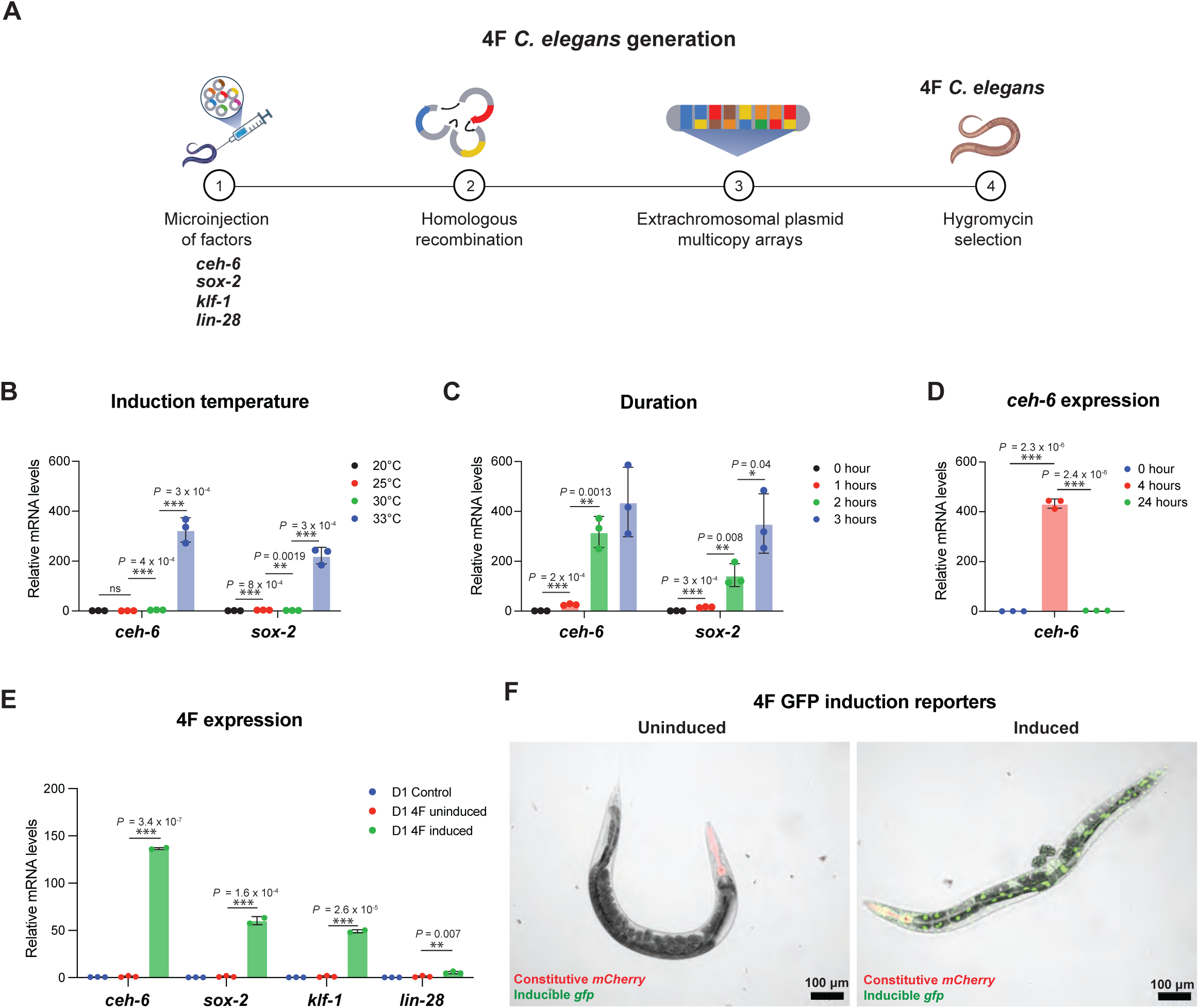
Generation of reprogrammable *C. elegans*. **A)** Schematic representation of the generation of 4F *C. elegans.* **B-C)** Relative mRNA levels of *ceh-6* and *sox-2* on day 1 (D1) induced 4F worms after heat-shock induction at different temperatures **(B)** and at different durations of 33°C heat-shock **(C)**. **D)** Relative mRNA levels of *ceh-6* on day 1 induced 4F worms at different times post-induction at 33°C for 3 hours. **E)** Relative mRNA levels of the four reprogramming factors on day 1 (D1) induced control, D1 uninduced 4F, and D1 induced 4F worms with an optimized induction protocol of heat shocking at 33°C for 3 hours. **F)** Representative confocal microscopic images of uninduced and induced D1 4F worms four hours post-induction at 33°C for 3 hours. Inducible GFP was used as a marker of 4F induction, while mCherry defined a constitutive marker indicating the presence of all 4 factors. Data show mean ± SEM. An ordinary two-way ANOVA test determined statistical significance. n=3 for all q-PCR samples.

### Reprogramming at different developmental stages causes morphological abnormality and premature death

To identify the effect of reprogramming during different developmental stages of *C. elegans*, reprogramming was induced at different developmental stages (Figure 2A). First, the induction of reprogramming in embryos resulted in embryos that were non-viable, with a significant 80% reduction in survival compared to the control group (Figure 2B and Figure 2C). Next, we induced in vivo reprogramming in L2 larval worms, observing morphological abnormalities as well as the inability to develop into adults (Figure 2D). Further characterization of these morphological defects showed a significant reduction of 60% in body size (Figure 2E), together with a significant reduction in median lifespan (Figure 2F). Lastly, the induction of in vivo reprogramming in L4 larval worms induced morphological alterations such as bagging (eggs retained inside the parental body) and internal hatching (eggs hatched inside the parental body) (Figure 2G), reduction in size (Figure 2H), and a significant reduction in the survival rate compared to control-induced worms (Figure 2I). Altogether, these results demonstrate that the induction of in vivo reprogramming during different developmental stages leads to high toxicity and developmental abnormalities, ultimately compromising the survival of worms.

**Figure 2:**
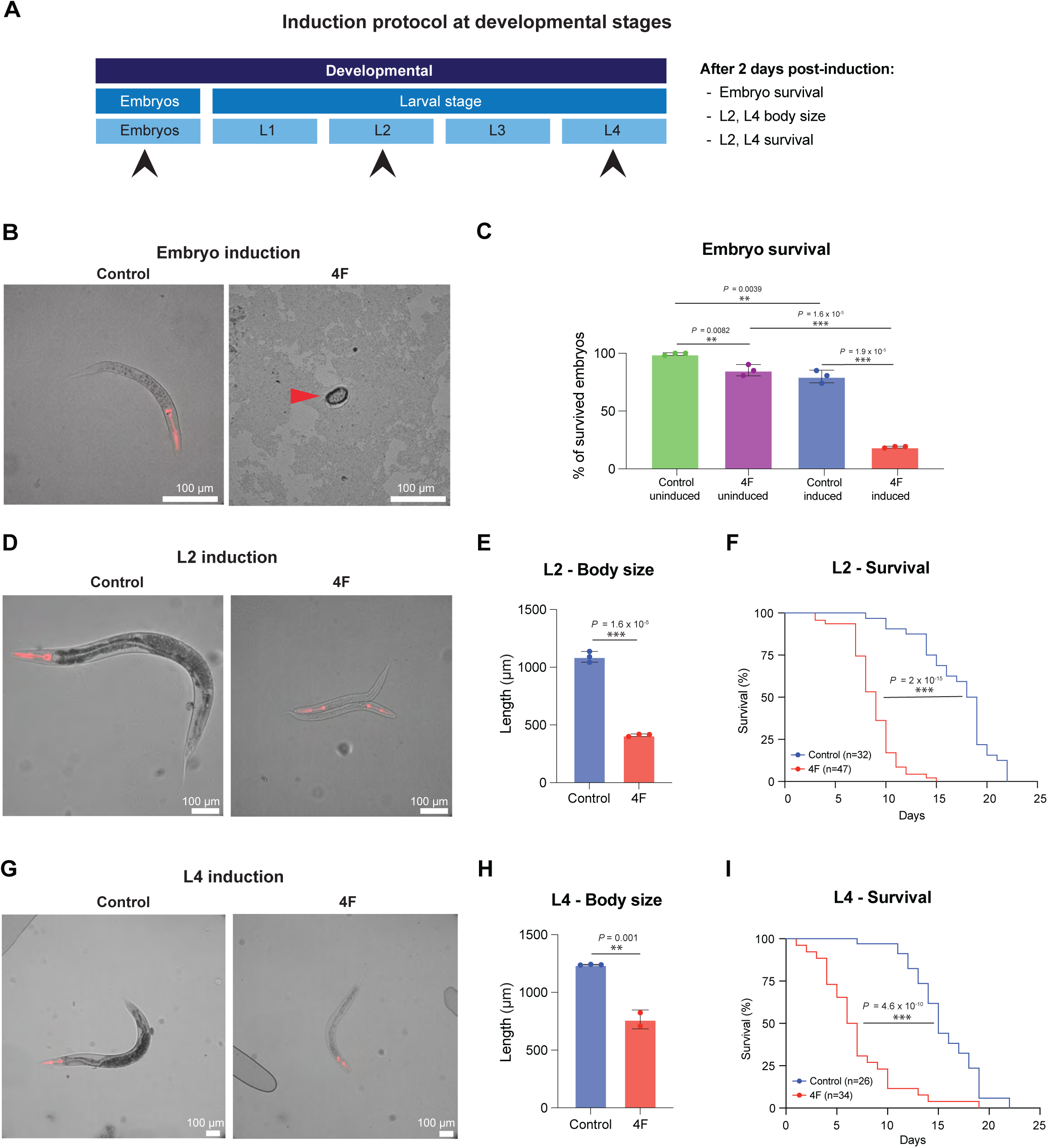
Reprogramming during developmental stages causes morphological abnormalities and premature death. **A)** Schematic representation of *C. elegans* developmental stages from embryos to larval L1, L2, L3, and L4 stage. Arrowheads indicate the developmental stages selected for induction. **B)** Representative confocal microscopic images of control and 4F worms were taken two days after induction at embryonic stage. The red arrow points to nonviable embryos. **C)** Analysis of survival of control and 4F worms two days after induction at embryonic stage. **D)** Representative confocal microscopic images of control and 4F worms two days post-induction at L2 larval stage. **E)** Length measurement of control and 4F worms two days post-induction at L2 stage. **F)** Analysis of survival of control and 4F worms after induction of reprogramming at the L2 stage. **G)** Representative confocal microscopic images of control and 4F worms two days post-induction at L4 larval stage. **H)** Length measurement of control and 4F worms at two days post-induction at L4 stage. **I)** Analysis of survival of control and 4F worms after induction of reprogramming at L4 stage. Data show mean ± SEM. An ordinary two-way ANOVA test determined statistical significance. n=3, for body size measurement.

### Reprogramming at reproductive stages causes morphological and behavioral abnormalities leading to premature death

Since the induction of reprogramming during development led to extreme toxicity, we decided to induce in vivo reprogramming in *C. elegans* during post-developmental stages. Towards this goal, we first induced the factors in young adults of age day 1 (D1), a time when worms are reproductively active^27^, and followed them for several days post-induction (Figure 3A). Importantly, D1 worms showed morphological abnormalities such as a significant size reduction and increased bagging upon induction of in vivo reprogramming (Figure 3B-D, Figure S2A). In addition, a significant reduction in egg-laying was observed in reprogrammed worms compared to their controls (Figure 3E). Moreover, behavioral abnormalities, including food avoidance (Figure 3F) and decreased motility, were also observed (Figure 3G). Finally, we observed a significant reduction in the survival rate following the expression of the reprogramming factors compared to control worms (Figure 3H).

**Figure 3:**
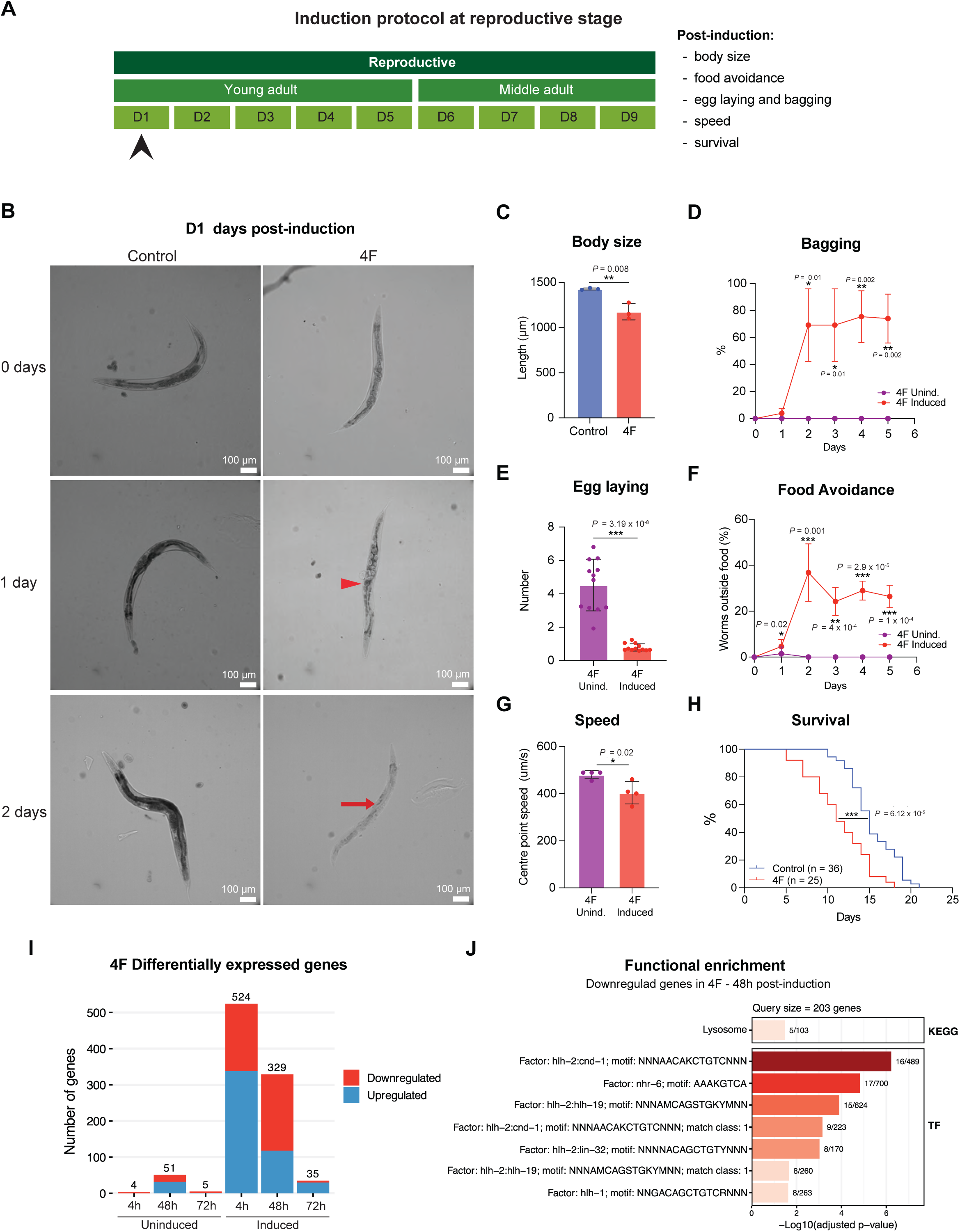
Induction of in vivo reprogramming during reproductive stages leads to morphological defects and premature death. **A)** Schematic representation of *C. elegans* reproductive stages. Arrowheads indicate the reproductive stage selected for induction. **B)** Representative confocal microscopic images of *C. elegans* control and 4F worms after induction at day 1 (D1) stage, on the day of induction (0 days), one day post-induction (1 day), and two days post-induction (2 days). Red arrowhead indicates bagging. Red arrow points to internal hatching. **C)** Length of control and 4F worms two days post-induction at D1. **D)** Percentage of bagging in 4F uninduced and induced worms over several days post-induction at D1. **E)** Analysis of number of eggs laid per worms of 4F uninduced and induced worms at two days post-induction at D1. **F)** Percentage of food avoidance of 4F uninduced (purple) and 4F induced (red) over several days post-induction at D1. **G)** Analysis of motility of 4F uninduced and induced worms measured two days post-induction at D1. **H)** Analysis of survival of control and 4F worms after induction on D1. **I)** Quantification of differentially expressed genes in 4F compared to the controls in the uninduced and induced at 4 hours, 48 hours, and 72 hours post-induction. **J)** Functional enrichment analysis of downregulated genes in 4F worms 48 hours post induction at D1. Data show mean ± SEM. An ordinary two-way ANOVA test determined statistical significance.

Next, in order to gain insight into the effects of in vivo reprogramming in *C. elegans*, we performed global transcriptome analysis by bulk RNA-sequencing. Analysis of RNA-seq showed a higher number of differentially expressed genes (DEGs) at 4 hours post-induction that were reduced over time at 48 and 72 hours post-induction (Figure 3I). Gene ontology (GO) analysis showed upregulation of genes related to sensory perception in 4F reprogrammable worms, which could explain food avoidance behavior (Figure S3A), and downregulation of genes related to immune response (Figure S3B).

In addition, functional enrichment analysis showed DEGs enrichment in transcription factor motifs related to development, such as *hlh-2*, *cnd-1*, *lin-14*, *che-1*, and *elt-3* (Figure 3J and Figure S3C-D). Altogether, these results demonstrated that the induction of reprogramming factors during post-developmental stages leads to toxicity and morphological abnormalities, ultimately compromising the survival of worms as well as the expression of developmental genes.

### Reprogramming of adult worms causes loss of proliferation and germ cell identity in the embryos and increased apoptosis without affecting somatic cell identity

Since the induction of in vivo reprogramming in D1 worms affected their reproductive capacity, we decided to focus on the study of embryos and the germline. First, we investigated the tissues where the reprogramming factors were expressed using the heat-inducible GFP marker. Upon inducing a D1 adult worm, we detected GFP expression corresponding to the four-factor expression in the embryos (Figure S4A). Importantly, no GFP expression was detected in the germline of the adult worm (Figure S4B). In addition, GFP was also detected in all major somatic tissues, such as head ganglions, intestine, and body wall muscle (Figure S4C-E). Next, to further characterize the effects of reprogramming at the cellular level, we crossed 4F non-GFP hermaphrodite worms with male worms carrying reporters for proliferation, germ cell, apoptosis, intestine, body wall muscle, and somatic cell identity (Figure S4F).

To test the effect of 4F induction on the proliferation rate, we induced reprogramming on 4F worms carrying a GFP proliferation reporter (4F.prol) on D1. Subsequently, we monitored the GFP signal for several days post-induction and observed a significant loss of proliferation at day 2 post-induction (Figure 4A-B), suggesting that 4F induction inhibits the proliferation of the embryos in adult worms. In order to further understand the effects of reprogramming in the embryos, we generated 4F worms with GFP germ cell reporter (4F.germ) to identify whether 4F induction could lead to an increase in germ cells. Subsequently, we induced the expression of the factors on the D1 4F.germ worms and detected the loss of germ cell identity in the embryos upon 2 days of reprogramming induction (Figure 4C). In addition, quantification of the germ cell identity signal showed a significant loss in the embryos of adult *C. elegans* upon reprogramming compared to their induced controls (Figure 4D).

**Figure 4:**
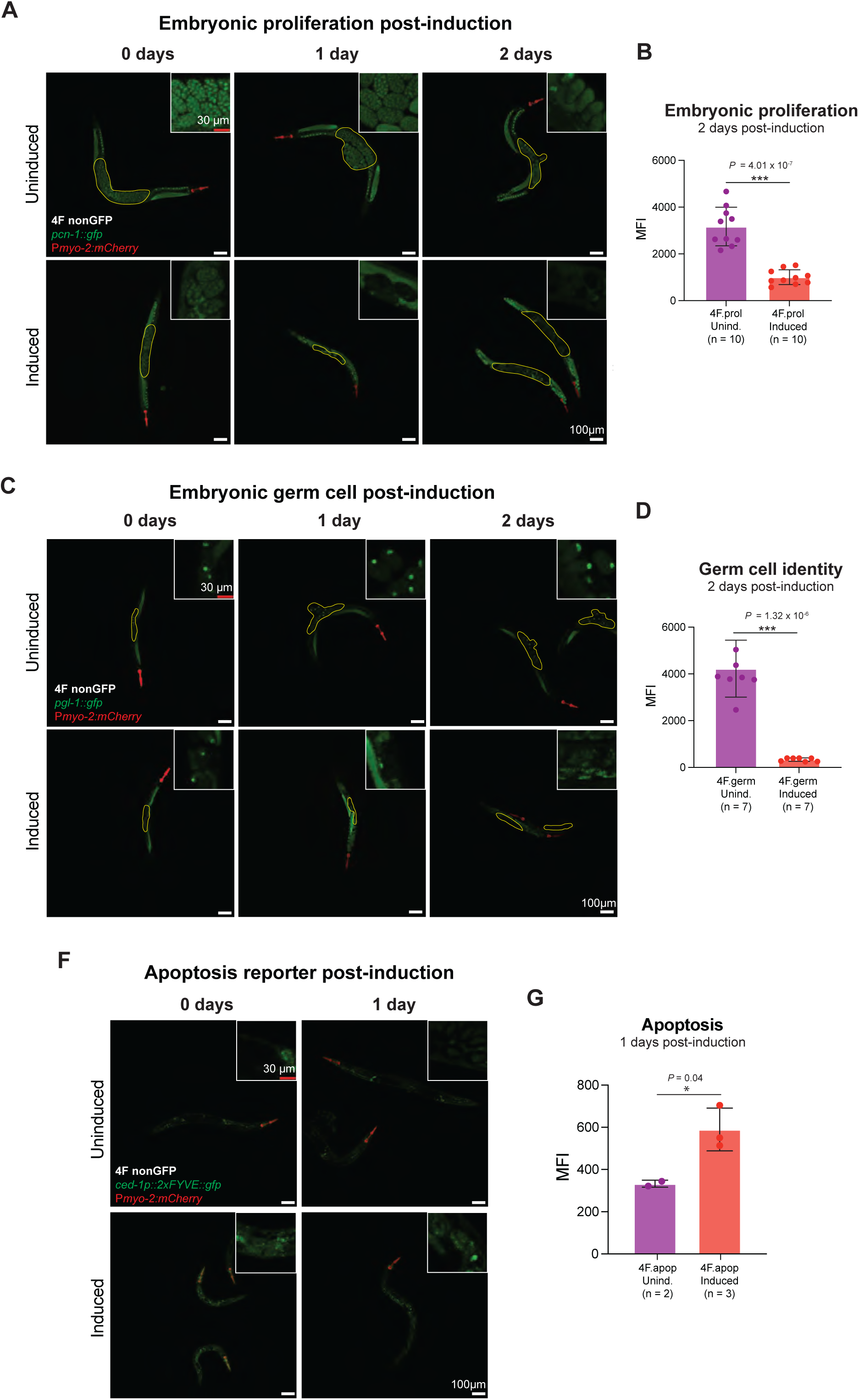
In vivo reprogramming at reproductive stages leads to loss of embryonic proliferation and germ cell identity together with increased apoptosis. **A**) Representative fluorescent images of 4F worms uninduced (top panel) and induced (bottom panel) at D1 stage, carrying the GFP proliferation reporter. Images were acquired on the day of induction (0 days) and 1- and 2 days post-induction. The yellow outline represents the region of interest for the GFP reporter within the worm. **B**) Quantification of embryonic proliferation in 4F worms uninduced and induced 2 days post-induction at D1 stage. **C**) Confocal fluorescent microscopy images of 4F worms uninduced (top panel) and induced (bottom panel) at D1 stage, carrying the GFP germ cell reporter, on the day of induction and 1- and 2-days post-induction. The yellow outline represents the region of embryos within the worms. **D**) Quantification of germ cell reporter in 4F worms 2 days post-induction at day 1 stage. **F**) Confocal fluorescent microscopy images of 4F worms uninduced (top panel) and induced (bottom panel) at D1 stage, carrying the GFP apoptotic reporter, at the day of induction and 1-day post-induction. **G**) Quantification of apoptotic GFP reporter fluorescence in 4F worms 2 days post-induction at day 1 stage. The white box represents a zoomed in portion of the region of interest. MFI: Mean fluorescent intensity. Data show mean ± SEM. An ordinary two-way ANOVA test determined statistical significance.

Since reprogramming could also lead to loss of cell identity and apoptosis^28^, we generated 4F worms carrying a GFP apoptotic reporter (4F.apop) to test the effect of 4F induction on the apoptosis rate. Next, we induced the reprogramming factors at D1 in the 4F.apop worms and observed a significant increase in apoptosis (Figure 4F and Figure 4G). In addition, to investigate whether reprogramming could also affect somatic tissues, we generated 4F worms with GFP intestinal reporter (4F.inte), TOM20 body wall muscle reporter (4F.bwm), and wrmScarlet somatic reporters (4F.soma). Subsequently, we induced reprogramming in these worms and monitored them for several days post-induction (Figure S4G-I). Importantly, compared to induced control worms, we did not detect significant differences in reporter signal associated with either the intestine, body wall muscle, or somatic tissue, suggesting that reprogramming did not affect the intestine, body wall muscle, or somatic cell identity (Figure S4J-L). Altogether, these results indicated that in vivo reprogramming of *C. elegans* leads to loss of proliferation and germ cell identity in the embryos of adult worms together with an increase in apoptosis throughout the body, while cell identity of post-mitotic tissues remains unaffected.

### Cyclic induction of reprogramming at post-reproductive stages leads to mild toxicity

As induction of in vivo reprogramming results in severe side effects during developmental stages as well as in the embryos of young adults, we decided to induce the expression of the reprogramming factors at post-reproductive stages. Towards this goal, *C. elegans* were subjected to heat shock at young adult on day 5 (D5), middle-aged adult on day 10 (D10), and old adult on day 15 (D15) (Figure 5A). Interestingly, the induction of reprogramming factors led to a 13% reduction in median lifespan at D5 and had no significant effects on survival at D10 and D15 compared to the induced controls (Figure 5B). Importantly, analysis of mRNA expression of the reprogramming factors at D10 showed a significantly lower expression of the factors compared to D1 (Figure 5C). In addition, and similar to D1, bulk RNA-seq analysis showed that the level of expression of the reprogramming factors was higher 4 hours post-inductions and decreased after 48 hours post-induction (Figure S5A). A similar trend was observed for DEGs, with a significantly lower number of DEGs in D10 compared to D1-induced worms 4 hours post-induction (Figure S5B). For this reason, we decided to test the effect of cyclic protocols for the induction of the reprogramming factors. Cyclic induction every 3 days or every 2 days did not have a significant impact on the survival rate compared to the induced controls. However, cyclic induction every day resulted in mild toxicity compared to uninduced controls, with a 10% reduction in median lifespan (Figure 5D). Together, these results suggest that *C. elegans* post-reproductive stages might be more resistant to the toxic effects of in vivo reprogramming than reproductive and developmental stages due to their lower plasticity and post-mitotic nature.

**Figure 5:**
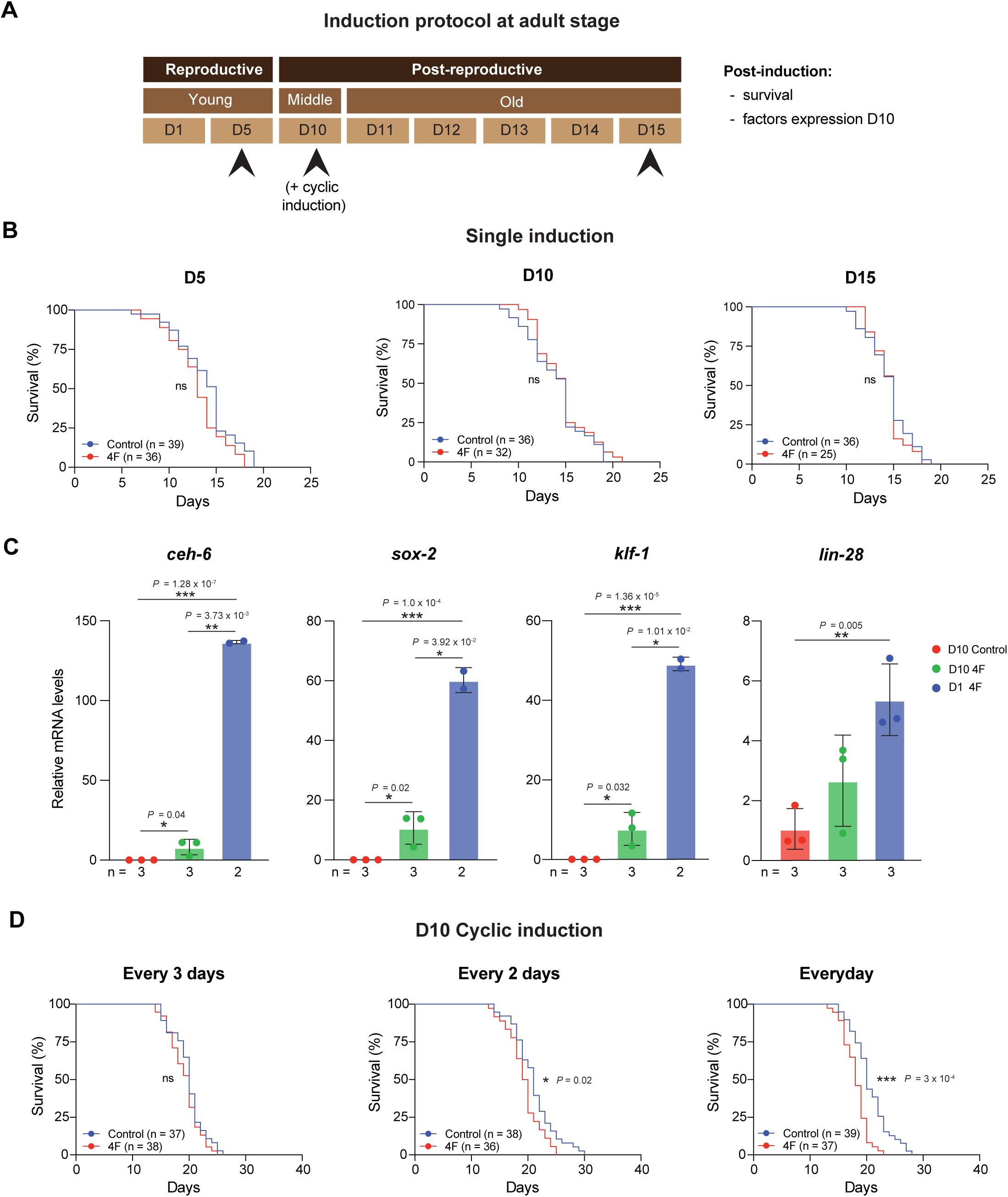
Reprogramming by cyclic induction of the reprogramming factors at post-reproductive stages results in mild toxicity. **A)** Schematic representation of post-reproductive stages of *C. elegans*. Arrowheads indicate the post-reproductive stages selected for induction. **B)** Analysis of survival of controls and 4F worms induced at day 5 (D5), day 10 (D10), and day 15 (D15) with single-shot induction. **C)** Relative mRNA expression levels of *ceh-6*, *sox-2*, *klf-1*, and *lin-28* in controls and 4F worms induced at D1 and D10. **D)** Analysis of survival of control and 4F worms induced at D10 stage with several cyclic induction protocols, once every three days, every 2 days, and every day. Data show mean ± SEM. An ordinary two-way ANOVA test determined statistical significance.

## DISCUSSION

### *C. elegans* as organism to study in vivo reprogramming

Cellular reprogramming has become increasingly important to investigate the interplay between epigenetics, cellular identity, and various biological processes, including regeneration and aging, yet numerous questions remain to be answered. Along this line, induction of in vivo reprogramming in mice is toxic and leads to organ dysfunction and tumor development resulting in premature death^28^. To date, no other organism has been used to study 4F in vivo reprogramming, raising the question of whether the effects observed in the mouse model could be similar in another organism. Importantly, *C. elegans* has been widely used as a model organism for research on development and aging. Here, we generated, for the first time, reprogrammable 4F worms to study the effects of in vivo reprogramming in vivo. The novel transgenic 4F *C. elegans* strains generated in this study allow the efficient induction of the reprogramming factors in adult somatic tissues, larvae, and embryos.

### Reprogramming at different developmental stages induces heterogeneous toxicity

Importantly, the induction of reprogramming factors at different stages of *C. elegans* development led to different degrees of toxicity. Specifically, reprogrammed embryos were non-viable, while reprogrammed L2s were unable to develop into reproductively active adults. Similarly, reprogrammed L4s became reproductive adults but showed bagging phenotype and premature death. In addition, L2 reprogramming worms displayed a stronger reduction in size compared to L4 and a more significant reduction in survival. These results demonstrate milder toxicity for the induction of in vivo reprogramming at advanced stages of development.

### Reprogramming at reproductive stages leads to bagging and food avoidance

During reproductive stages, we observed GFP expression corresponding to the 4F expression in the developing embryos, but no expression was achieved in the germline. Although germline expression at low levels can be achieved from single-copy transgenes (REF), transgene arrays30, including hsp16 promoter-bearing ones31, have been previously shown to be vulnerable to silencing in the germline.

This might be due to the tight epigenetic regulation, which has been reported and is linked to the silencing of transgenes in the germline^29^. Although germline expression at low levels can be achieved from single-copy transgenes^30^, transgene arrays^31^, including hsp16 promoter-bearing ones^32^, have been previously shown to be vulnerable to silencing in the germline. Reprogramming of *C. elegans* at D1 led to strong morphological and behavioral abnormalities such as bagging, reduced egg laying, reduced motility, and food avoidance. Interestingly, the RNA-seq analysis showed downregulation of genes related to vulval *hlh-2* and embryonic development *cnd-1*, potentially explaining the bagging and dysfunctional embryonic development. In addition, genes related to sensory perception were upregulated in 4F reprogrammable worms, potentially explaining food avoidance behavior.

### Reprogramming at reproductive stages adversely affects the developing embryos

Previous studies have demonstrated that cellular reprogramming of mammalian cells induces cell proliferation and leads to the expression of pluripotency markers^1^. However, due to the lack of a pluripotency state in adult *C. elegans*, we focused on the study of both germline and proliferation reporters. Surprisingly, reprogramming in *C. elegans* resulted in the loss of germ cell identity and a decrease in proliferation in developing embryos in the adult worms. In addition, functional enrichment analysis showed DEGs enrichment in transcription factors related to development. One of the transcription factor motifs that was upregulated was *lin-14*, a temporal regulator of postembryonic developmental events. Importantly, this gene is only expressed until the L2 stage during normal development. Another transcription factor motif that was upregulated is zinc finger transcription factor *che-1*, which has been previously used for reprogramming germ cells into neurons^24^. Interestingly, the expression of the GATA transcription factor motif *elt-3* was also affected. Importantly, previous studies have shown that the ectopic expression of GATA transcription factor *elt-7* can reprogram the pharynx or somatic gonad into intestinal fate^25^. Our findings highlight the potential of in vivo reprogramming to influence cellular plasticity in *C. elegans*. Nevertheless, somatic cell identity remained largely unaffected upon induction of reprogramming at later adult stages, likely due to the limited plasticity of the post-mitotic cell stage.

### Reduced toxicity of reprogramming at post-reproductive stages following a cyclic induction protocol

Induction of in vivo reprogramming at post-reproductive stages, such as day 5, 10, and 15 worms, did not affect survival. This could be mainly attributed to two reasons: 1) lower expression levels of the reprogramming factors or 2) absence of embryos at these post-reproductive stages. Moreover, the absence of embryos could explain the lower levels of expression at D10 compared to D1 induction. In addition, RNA analysis on D10 worms showed a lower number of DEGs compared to D1-induced worms. However, the cyclic induction of reprogramming factors every day led to a mild reduction in the median lifespan, which could be caused by the continuous expression of the reprogramming factors in the somatic tissues of adult worms.

In summary, these results demonstrate that the induction of in vivo reprogramming results in different degrees of toxicity that are inversely correlated with development and, most likely, cellular plasticity. Overall, this study represents the first attempt to induce in vivo reprogramming by expressing Yamanaka-like factors in *C. elegans*. Like previous studies in mice, induction of in vivo reprogramming is highly toxic and lethal, especially during development. Using invertebrate model organisms such as *C. elegans* to study in vivo reprogramming might increase our understanding of the interplay between epigenetics and cellular identity during development, adulthood, and aging.

### Limitations of the study

Induction of in vivo reprogramming predominantly affected embryos, larvae, and reproductively active young adults, where cells are characterized by a higher degree of plasticity. Nevertheless, no expression was achieved in the germline, possibly due to the silencing of transgenes in the germline. In addition, we are aware that the post-mitotic nature of *C. elegans* cells represents a major limitation for studying cellular reprogramming in this organism. In this line, previous studies have shown that some epigenetic factors can hinder somatic cell reprogramming^17,18^. Further studies focused on removing these epigenetic barriers in combination with the expression of the reprogramming factors might allow a higher degree of cellular plasticity upon induction of cellular reprogramming in adult *C. elegans*.

Importantly, the generation of 4F worms using CRISPR integration led to high lethality, most probably due to the leakiness or induction of the reprogramming factors during the process of genetic modification and their high toxicity during development. For this reason, an extrachromosomal multi-copy array, where the copy number of factors cannot be controlled, was used to generate reprogramming *C. elegans* strains. Lastly, despite the common use of heat shock promoters in *C. elegans*, heat shock itself can have toxic effects and be lethal beyond a certain temperature or duration.

Multiple studies have previously demonstrated the induction and need for cellular proliferation during cellular reprogramming to pluripotency. Since our data does not demonstrate complete reprogramming of *C. elegans* cells to stemness or pluripotency, future experiments, including analysis of tissue-specific or cell-specific transcriptional changes, will be necessary to understand better the effect of in vivo reprogramming on cellular identity. Moreover, tissue-specific reprogramming could also be a better way to understand this^33^. This analysis might provide a better understanding of the differences in toxicity observed upon induction of in vivo reprogramming at the D1 and D10 stages. Moreover, germline-less 4F worms lacking germ cells will allow us to dissect the role of germ cells and embryos on the toxic effect of the reprogramming factors in young adults.

## METHODS

### Generation of transgenic 4F strains

To generate reprogrammable *C. elegans*, orthologs of the murine reprogramming factors *Oct4*, *Sox2*, *Klf4,* and *Lin28* were identified using bioinformatic analysis. The corresponding orthologs *ceh-6*, *sox-2*, *klf-1*, and *lin-28 in C. elegans* were codon-optimized (Supplementary Table 1). These factors were cloned under the control of a heat-shock (*hsp-16.2*) promoter^34^. In addition to all four factors, a heat-inducible GFP marker under the *hsp-16.2* promoter, a constitutive mCherry marker under the *myo-2* promoter, and a hygromycin resistance selection marker were also cloned (Figure S1A). Subsequently, reprogramming factors *ceh-6, sox-2, klf-1, and lin-28* were co-injected with selection markers into the *C. elegans* germline resulting in the formation of extrachromosomal multicopy array by homologous recombination of the plasmids, and a final selection of hygromycin resistant 4F transgenic worms (Supplementary Table 2). Finally, three types of transgenic worms were generated: a reprogrammable *C. elegans* strain containing the four factors with an inducible GFP marker as 4F, a strain lacking the four factors that serve as a control, and a strain without an inducible GFP marker as 4F non-GFP.

### Induction protocols

The standard induction protocol for reprogramming factors in 4F *C. elegans* involved incubating the embryos or worms in an incubator (Memmert) for a 3-hour heat shock of 33°C or the duration and temperatures indicated in the manuscript. Following the induction, *C. elegans* was moved back to the maintenance temperature of 20°C.

### Generation of 4F reporter strains

The reporter strains used in this study, DQM662 (Proliferation), JH3269 (Germline), ZH231 (Apoptosis), SJ4143 (Intestine), PS6192 (Body wall muscle), and WBM136 (Somatic), were obtained from the Caenorhabditis Genetic Center (CGC). To generate 4F reporter strains, the 4F non-GFP, carrying mCherry reporter strain, was crossed with each of the above-mentioned reporters. Briefly, male progeny reporters were generated by heat shocking L4 stage larvae at 30°C for 6 hours. For the crossing, a total of 5 L4 stage hermaphrodite 4F non-GFP worms were placed in a P60 NGM plate with 5 young male reporter worms. Worms showing both reporters were selected and maintained (Supplementary Table 3).

### Survival experiment

Worms were grown on solid NGM plates (P60 plate, Falcon, 353004) seeded with UV-killed OP50 bacteria as food (150ul of 120 mg/ml UV killed bacteria per P60 plate with 10 ml NGM) at 20°C. Hygromycin selection plates were seeded with 4 mg/mL hygromycin B (Hygromycin B Gold, InvivoGen, HGG-44-04). Worm populations were synchronized by hypochlorite treatment (Bleach solution:2.8% bleach, 0.8N NaOH). Pelleted worms were treated with bleach solution for up to 12 minutes until all the worms rapture and eggs were released. Immediately, eggs were washed with ddH20 up to 2 times. The washed eggs were seeded onto an NGM plate without food for 24 hours. After seeding OP50 bacteria, we waited until the worm reached the L4 stage. For survival experiments, L4 worms were sterilized by 5-fluorodeoxyuridine treatment (CAS number 50-91-9, Acros organics), and lifespan was measured by testing the worm movement with gentle poking using a worm pick. Worms were counted as dead when they showed no movement at all.

### Confocal Microscopy

For confocal image acquisitions, worms were mounted on a fresh 4% low-melting agarose (NuSieve^TM^ GTG^TM^ Agarose, Lonza, 50081) pad between glass slides (Epredia, Superfrost Plus^TM^ Gold Adhesion Microscope Slides, K5800AMNZ72). First, 10-15 worms were individually picked and transferred into an agar layer containing 10 mM levamisole, used as a paralyzing agent. All the confocal microscopic worm images were captured using a Nikon Ti2 Yokogawa CSU-W1 spinning disk confocal microscope with NIS Elements software. The images were analyzed using FIJI Version: 2.14.0/1.54f.

### Behavioral analysis

The food avoidance test was done on synchronized 4F-induced worms by counting the number of worms that stayed out of the food compared to those that stayed within the food. The movement analysis was done by capturing a minute video of worms using a Nikon SMZ800N microscope with a coupled camera. The videos acquired were analyzed using MBF bioscience Worm Tracker.

### Morphological analysis

The morphological analysis and size measurements were performed on 10x images of the worms acquired with Nikon Ti2 Yokogawa CSU-W1 spinning disk confocal microscopy. Later, the quantification of worm length was measured using FIJI Version 2.14.0/1.54f. Bagging was measured by manual counting, by manually looking at worms that have bagging phenotypes, and those without using 2x objective in a stereo microscope Motic SMZ-171. The egg-laying rate was measured by acquiring images of NGM plates with the worms using a Motic SMZ-171 stereo microscope with an objective of 2x.

### Quantitative real-time PCR

Total RNA was extracted from synchronized worm populations by TRIzol reagent (Invitrogen, 15596018) and chloroform (Roth, 280298902) treatment. RNA was then purified using Monarch Total RNA miniprep Kit (New England Biolabs, T2010S) according to the manufacturer’s instructions. Samples were treated with DNase (Qiagen, 79254) for 15 minutes (1:8 in DNase buffer). Total RNA concentrations were determined using the Qubit RNA BR Assay Kit (Thermofisher, Q10211). cDNA synthesis was performed by adding 4 μL of iScript™ gDNA Clear cDNA Synthesis (Biorad, 1725035BUN) to 500ng of RNA sample and run in a Thermocycler (Biorad, 1861086) with the following protocol: 5 min at 25°C for priming, 20 min at 46°C for reverse transcription, and 1 min at 95°C for enzyme inactivation. Final cDNA was diluted 1:5 using autoclaved water and stored at −20°C. qRT-PCR was performed using SsoAdvanced SYBR Green Supermix (Bio-Rad, 1725272) in 384 well PCR plates (Thermofisher, AB1384) using the QuantStudio™ 12K Flex Real-time PCR System instrument (Thermofisher). Forward and reverse primers (1:1) were used at a final concentration of 5 µM with 1 µL of cDNA sample (Supplementary Table 4).

### RNA-seq alignment and quantification

Data was processed using nf-core/rnaseq v3.14.0 (doi: https://doi.org/10.5281/zenodo.1400710) of the nf-core collection of workflows ^35^, utilising reproducible software environments from the Bioconda^36^ and Biocontainers^37^ projects. The pipeline was executed with Nextflow v23.10.0^38^ with the following command:

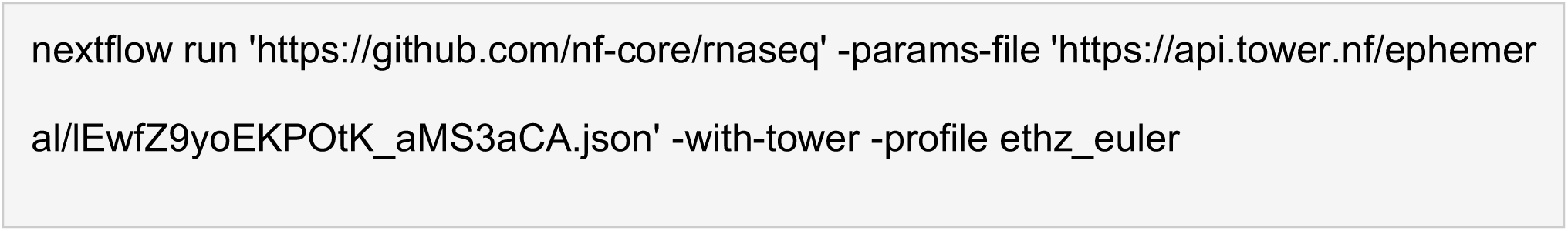

The RNA-seq reads were aligned to the *Caenorhabditis elegans* reference genome WBcel235 (GCA_000002985.3), and gene annotation was obtained from the Ensembl release 111.

To align and quantify the reads assigned to the extrachromosomal array sequence, the nf-core/rnaseq pipeline version 3.14.0 was used, having has input the unmapped reads generated from the pipeline run described above. Prior to alignment, rRNA contaminants were removed using the ‘SortMeRNA’ package^39^.

The pipeline was executed with Nextflow v23.10.1 with the following command:

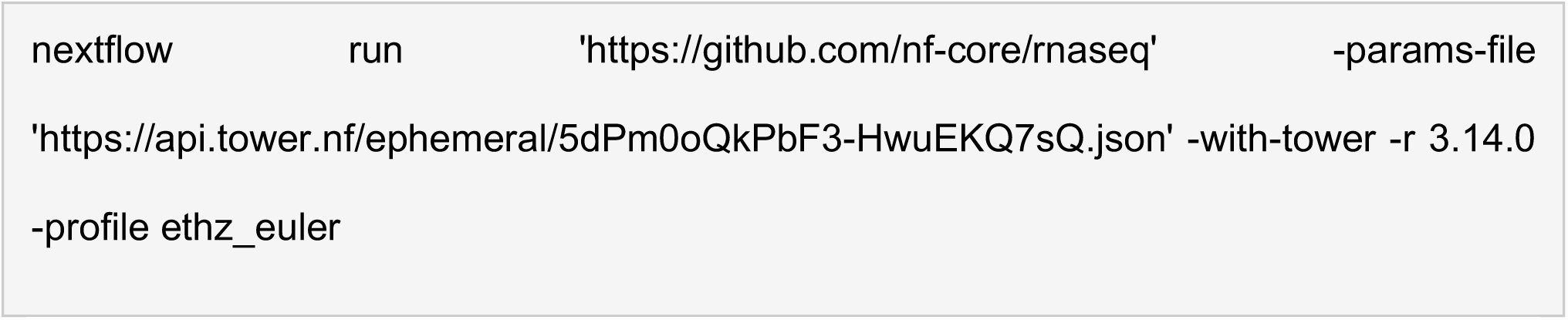

### RNA-seq analysis

Transcript read counts were imported into R and converted to gene counts using the Bioconductor package ‘tximport’ ^40^. Normalization was conducted using the Bioconductor package ‘DESeq2’^41^, and dimensionality reduction was performed using the reads counts after variance stabilizing transformation (VST).

Differential expression analysis was carried out using the ‘DESeq2’ package with the parameter modelMatrixType set to “standard” and default settings for other parameters. Genes were considered differentially expressed between conditions if they exhibited an adjusted p-value below 0.05 and an absolute log2 fold change exceeding 2.

Gene ontology analysis was conducted using the ‘compareCluster’ function from the Bioconductor package ‘clusterProfiler’^42^. The analysis utilized the org.Ce.eg.db database, focusing solely on Biological Process (BP) ontology. Benjamini-Hochberg adjustment was applied for p-values, with significance thresholds set to 0.05 for both p-values and q-values.

### Statistical analysis

Statistical analysis was performed using GraphPad Prism 9.4.1 (GraphPad Software). For the comparison of two independent groups, a two-tailed unpaired t-Student’s test (data with normal distribution) was executed. The corresponding p and n values are presented in the indicated figures, and the levels of significance are denoted as follows: ∗∗∗p < 0.001, ∗∗p < 0.01, ∗p < 0.05, and ns indicates not significant. n values represent the number of animals. Data shown mean ± standard error mean.

## ACKNOWLEDGMENTS

The authors thank all members of the Ocampo laboratory, especially Alba Vílchez-Acosta, Gabriela Desdín-Micó, and María del Carmen Maza, for their valuable feedback and support. In addition, we would also like to thank the UNIL Cellular Imaging Facility and Rosa Chiara Paolicelli (Assistant Professor, UNIL, Switzerland) and Anne-Claire Companion (Postdoc, UNIL, Switzerland) for their valuable support with microscopy and imaging. We thank Pierre Gönczy (Professor, EPFL, Switzerland) for his valuable support. Some strains were provided by the CGC, which is funded by the NIH Office of Research Infrastructure Programs (P40 OD010440).

## AUTHOR CONTRIBUTIONS

A.O. and N.K. designed the study. N.K. was involved in all experiments, data collection, analysis, and interpretation. A.V.A. and S.P. revised the manuscript and prepared the figures. Y.M. contributed to survival experiments, confocal microscopy, and qRT–PCR analysis under the supervision of N.K. M.P., C.F.J., and S.E.M. generated the transgenic 4F worms. V.P., F.v.M. and J.A.S performed bioinformatic analysis and analysis of RNA-seq data. A.O. directed and supervised the study and designed the experiments. N.K. and A.O wrote the manuscript with input from all authors.

---

**Figure S1:**
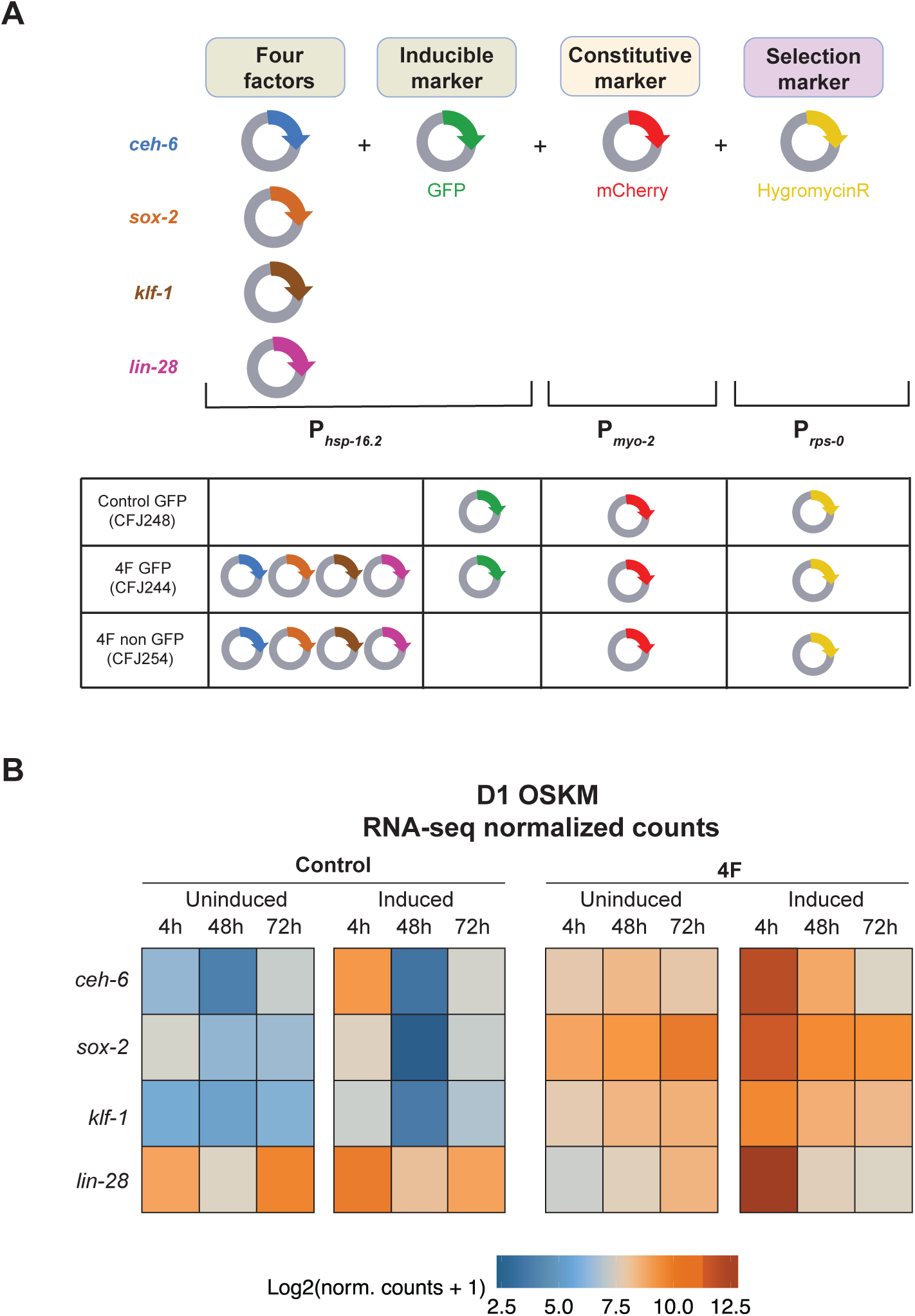
Generation of 4F *C. elegans* strains. **A)** Schematic representation of plasmid constructs used in the injection mixture for the generation of 4F worms. All four factors were cloned under the control of the heat shock promoter *hsp-16.2*. GFP was used as an inducible marker under the control of the *hsp-16.2* promoter, mCherry as a constitutive marker under the pharynx-specific *myo-2* promoter, and hygromycinR as a selection marker under the *rps-0* promoter. **B)** Heatmap from RNA-seq analysis showing the relative expression of *ceh-6*, *sox-2*, *klf-1*, and *lin-28* in control and induced and uninduced 4F worms 4 hours, 48 hours, and 72 hours post-induction at D1.

**Figure S2:**
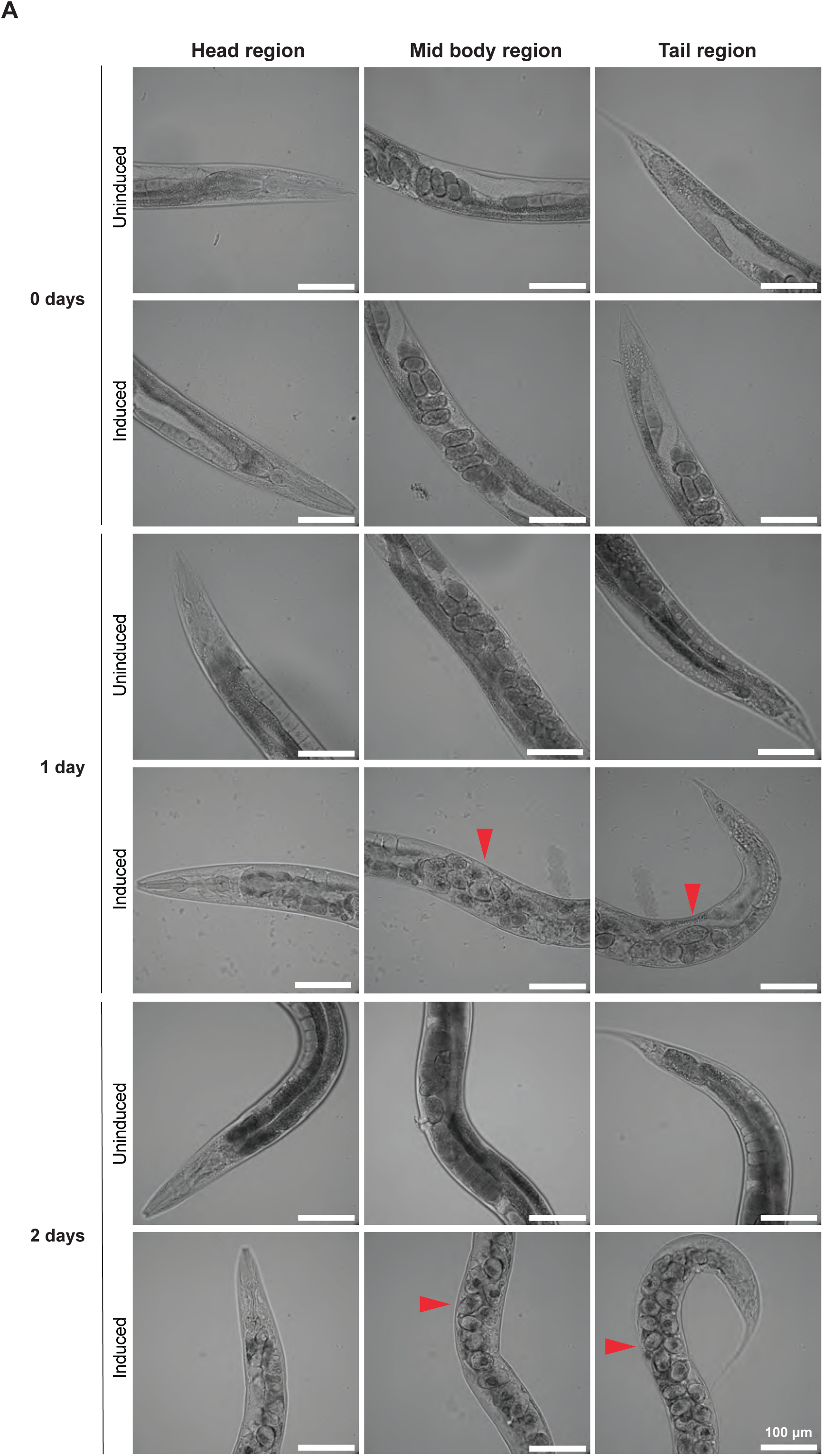
Bagging in D1-induced 4F worms. **A)** Representative confocal microscopic images of the head, midbody, and tail region of 4F D1 induced and uninduced worms at 0 days, 1 day, and 2 days post-induction. The arrowhead indicates the bagging phenotype. Scale bar 100 µm.

**Figure S3:**
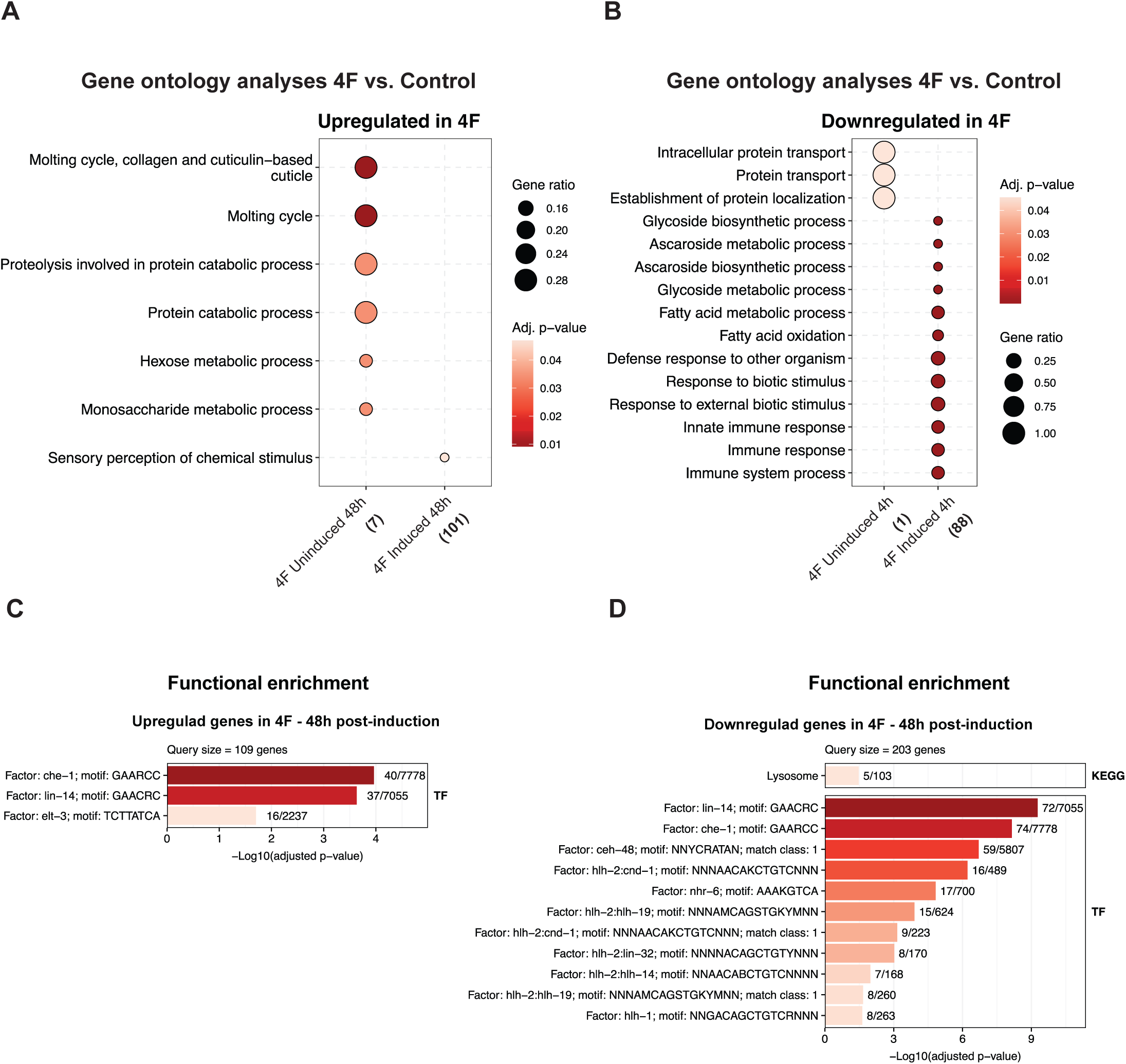
Upregulation of genes related to sensory perception and transcription factors *lin-14*, *che-1*, and *elt-3* in D1-induced 4F worms. **A-B)** Gene ontology analysis showing upregulated (A) and downregulated (B) pathways of 4F D1-induced and uninduced worms. **C-D)** Functional enrichment of upregulated (C) and downregulated (D) genes in 4F induced 48 hours post-induction.

**Figure S4:**
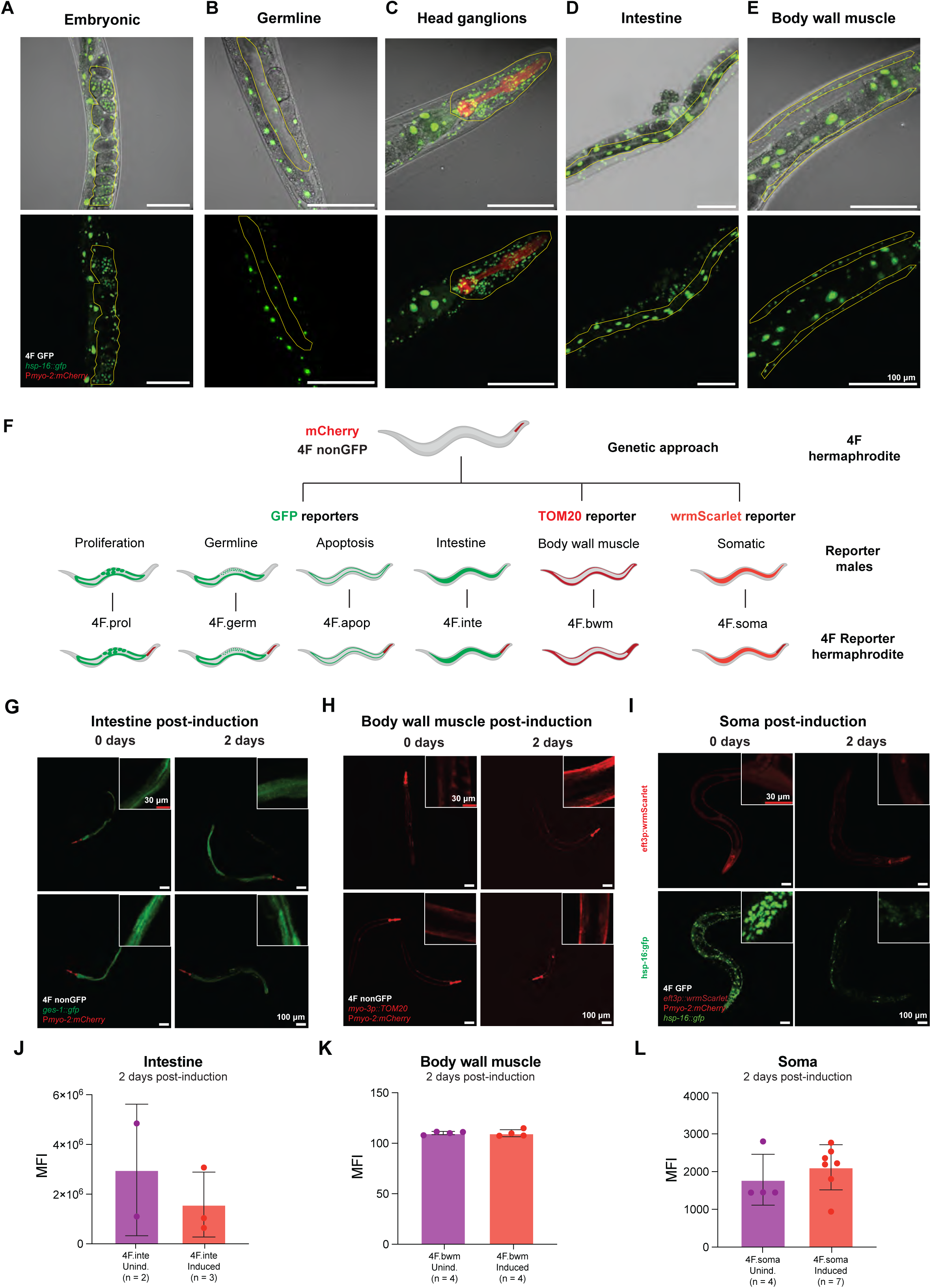
Preservation of cell identity in D1-induced 4F worms. **A-E)** Confocal fluorescent microscopy images of 4F worms induced at the D1 stage carrying the GFP reporter for heat-shock corresponding to the 4F expression in the embryos (A), germline (B), head ganglions (C), intestine (D), and body wall muscle (E), 4 hours post-induction. **F)** Schematic representation of the generation of 4F reporter worms by crossing 4F non-GFP hermaphrodites with males carrying fluorescent reporters for proliferation, germline, apoptosis, intestine, body wall muscle, and somatic cell identity. **G)** Confocal images of 4F GFP intestinal reporter worms (4F.inte) were imaged on 0d PI and 2d PI. The yellow outline represents the region of interest for the GFP reporter within the worm. **H)** Confocal images of TOM20 body wall muscle reporter worms (4F.bwm) imaged on 0d PI and 2d PI. **I)** Confocal images of wrmScarlet somatic reporter worms (4F.soma) imaged on 0d PI and 2d PI. The top panel indicates the somatic cells, and the bottom panel shows 4F expression on 0d PI and 2d PI. **J)** Quantification of confocal fluorescent images of 4F.inte-HS and 4F.inte+HS on 2d PI. **K)** Quantification of confocal fluorescent images of 4F.bwm-HS and 4F.bwm+HS on 2d PI. **L)** Quantification of confocal fluorescent images of 4F.soma-HS and 4F.soma+HS on 2d PI. The white box represents a zoomed portion of the region of interest. MFI: Mean fluorescent intensity. Data show mean ± standard mean. An ordinary two-way ANOVA test determined statistical significance.

**Figure S5:**
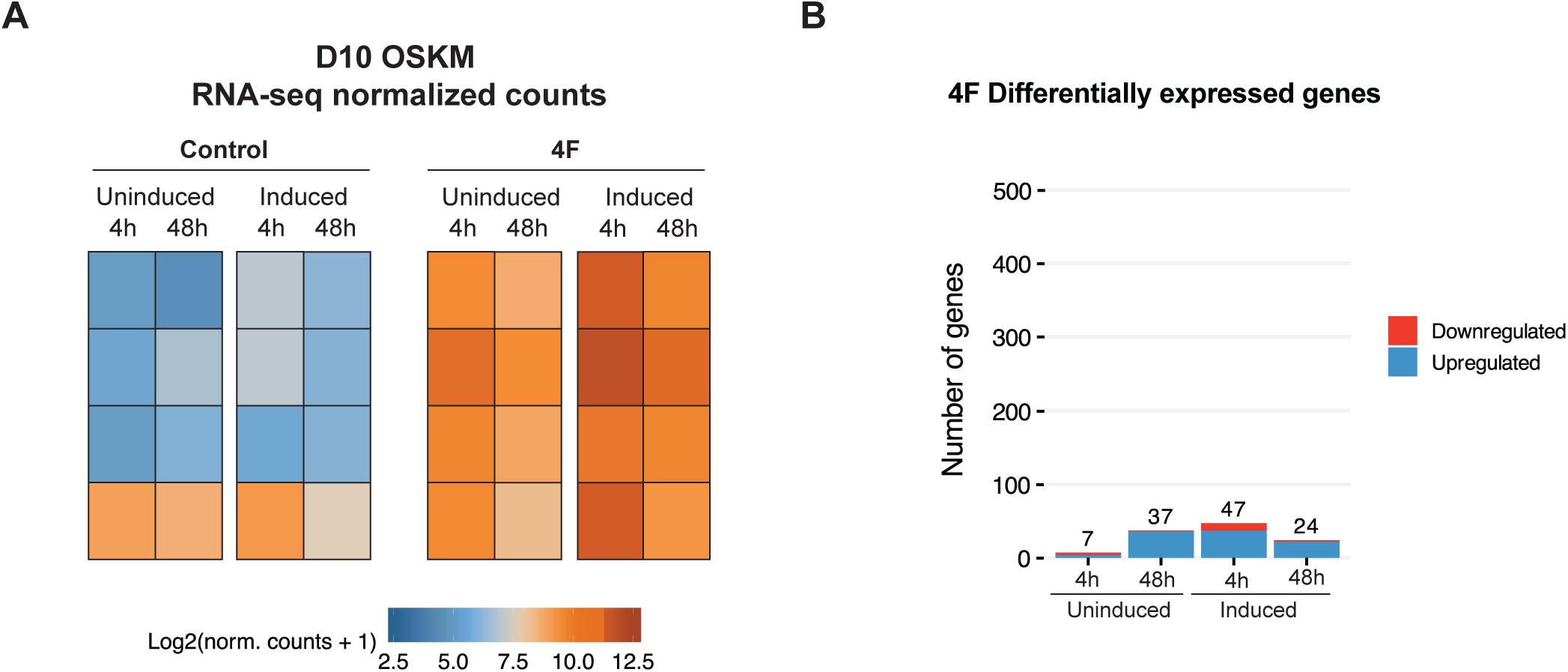
Decreased number of differentially expressed genes in 4F worms induced at D10. **A)** Heatmap from RNA-seq analysis showing the relative expression of *ceh-6*, *sox-2*, *klf-1*, and *lin-28* in controls and 4F induced and uninduced worms 4 hours and 48 hours post-induction at day 10. **B)** Analysis of differentially expressed genes in 4F uninduced and induced worms 4-hour and 48-hour post-induction at day 10 worms.

