## Supplementary Table 1 for "In vivo reprogramming of *Caenorhabditis elegans* leads to heterogeneous effects on lifespan"

| cceh6_pPD49_78 | csox2_pPD49_78 | cklf1_pPD49_78 | clin28_pPD49_78 |
|---|---|---|---|
| Sequence | Sequence | Sequence | Sequence |
| [illegible] | [illegible] | [illegible] | [illegible] |

### cceh6 pPD49_78 MAP

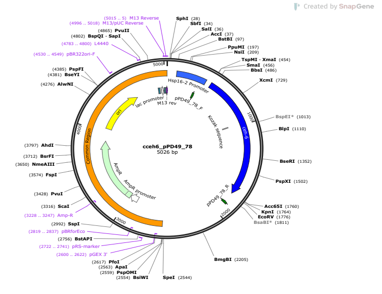


### csox2 pPD49_78 MAP

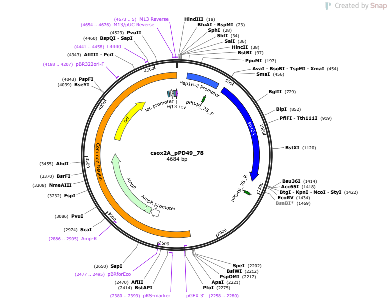


### cklf1 pPD49_78 MAP

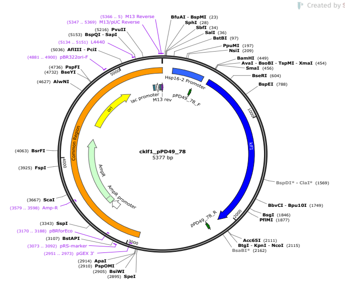


### clin28 pPD49_78 MAP

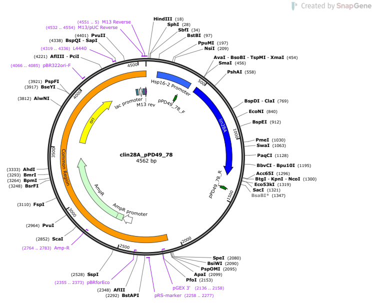
