## Supplementary Table 2 for "In vivo reprogramming of *Caenorhabditis elegans* leads to heterogeneous effects on lifespan"

| Label in manuscript | Strain | Genetic background | Allele | Transgenes | Injection |
|---|---|---|---|---|---|
| 4F | CFJ244 | N2 | kstEx77 | L4043 (Phsp-16.2::gfp) @ 5 ng/ul, pCFJ782 (HygroR) @ 15 ng/ul, pCFJ90 (pmyo-2::mCherry) @ 10 ng/ul, ce-ceh6 in pPD49.78 @ 5 ng/ul, ce-sox2A in pPD49.78 @ 5 ng/ul, ce-klf1 in pPD49.78 @ 5 ng/ul, ce-lin28A in pPD49.78 @ 5ng/ul, 1 kb plus Invitrogen ladder @ 50 ng/ul. | **iSEM883** |
| Control | CFJ248 | N2 | kstEx81 | L4043 (Phsp-16.2::gfp) @ 5 ng/ul, pCFJ782 (HygroR) @ 15 ng/ul, pCFJ90 (pmyo-2::mCherry) @ 10 ng/ul, 1 kb plus Invitrogen ladder @ 70 ng/ul. | **iSEM884** |
| 4F non GFP | CFJ254 | N2 | kstEx87 | pCFJ782 (HygroR) @ 15 ng/ul, pCFJ90 (pmyo-2::mCherry) @ 10 ng/ul, ce-ceh6 in pPD49.78 @ 5 ng/ul, ce-sox2A in pPD49.78 @ 5 ng/ul, ce-klf1 in pPD49.78 @ 5 ng/ul, ce-lin28A in pPD49.78 @ 5ng/ul, 1 kb plus Invitrogen ladder @ 55 ng/ul. | **iSEM886** |
