## Supplementary Table 3 for "In vivo reprogramming of *Caenorhabditis elegans* leads to heterogeneous effects on lifespan"

| No | Reporter strains | Reporter for | Genotype | Description | Ordering link (CGC) |
|---|---|---|---|---|---|
| 1 | DQM662 | Proliferation | bmdSi200 I; bmdSi168 II. | bmdSi200 [loxN::pcn-1p::pcn-1::GFP] I. bmdSi168 [loxN::rps-27p::DHB::2x-mKate2] II. | https://cgc.umn.edu/strain/DQM662 |
| 2 | JH3269 | Germcell | pgl-1(ax3122[pgl-1::gfp]) IV. | GFP inserted at C terminus of endogenous pgl-1 locus. | https://cgc.umn.edu/strain/JH3269 |
| 3 | ZH231 | Apoptotic | enIs7 X. | enIs7 [ced-1p::ced-1::GFP + unc-76(+)] X. | https://cgc.umn.edu/strain/ZH231 |
| 4 | PS6192 | Body wall muscle | syIs243. | syIs243 [myo-3p::TOM20::mRFP + unc-119(+) + pBS Sk+]. | https://cgc.umn.edu/strain/PS6192 |
| 5 | SJ4143 | Intestine | zcIs17. | zcIs17 [ges-1::GFP(mit)]. | https://cgc.umn.edu/strain/SJ4143 |
| 6 | WBM1364 | Somatic | wbmIs119 V. | wbmIs119 [eft-3p::3XFLAG::rpl-22::SL2::wrmScarlet::unc-54 3'UTR, *wbmIs88] V. | https://cgc.umn.edu/strain/WBM1364 |

| No | 4F Reporter strains | Label in manuscript |
|---|---|---|
| 1 | DQM662 CFJ254 | 4F.prol |
| 2 | JH3269 CFJ254 | 4F.germ |
| 3 | ZH231 CFJ254 | 4F.apop |
| 4 | PS6192 CFJ254 | 4F.bwm |
| 5 | SJ4143 CFJ254 | 4F.inte |
| 6 | WBM1364 CFJ254 | 4F.soma |
