## Supplementary Table 4 for "In vivo reprogramming of *Caenorhabditis elegans* leads to heterogeneous effects on lifespan"

| No | Gene | Forward | Reverse |
|---|---|---|---|
| 1 | *ceh-6* | CTCCGAGCCATCTTCTCTTAAC | CGTAGTTAGCAGCCTCCATTT |
| 2 | *sox-2* | CTAAGCGTCTTCGTGCTATCC | GATTGGAGCTCCGTTCTTCTT |
| 3 | *klf-1* | CTACCTCTTCTTCTTCCT | GTGAGAGTTCTTGTATGG |
| 4 | *lin-28* | TTCCTTCCTTCGAGAGGCTCC | TCCTTCATCAAGACTCCGAAATCC |
| 5 | *act-1* | ACGACGAGTCCGGCCCATCC | GAAAGCTGGTGGTGACGATGGTT |
| 6 | *tba-1* | TCAACACTGCCATCGCCGCC | TCCAAGCGAGACCAGGCTTCAG |
| 7 | *pmp-3* | TGGCCGGATGATGGTGTCGC | ACGAACAATGCCAAAGGCCAGC |
| 8 | *gfp* | TTCTGTTATGGTGTTCAATG | GTAGTTCCCGTCATCTTT |
| 9 | *mCherry* | CGGCAGATATACCAGATTA | GTCACTGTAACAACTCCT |
